# Dynamic FRET-FLIM based screens of signal transduction pathways: a feasibility study

**DOI:** 10.1101/2021.05.23.445328

**Authors:** Rolf Harkes, Olga Kukk, Sravasti Mukherjee, Jeffrey Klarenbeek, Bram van den Broek, Kees Jalink

## Abstract

Fluorescence Lifetime Imaging (FLIM) is an intrinsically quantitative method to screen for protein-protein interactions and frequently used to record the outcome of signal transduction events. With new highly sensitive and photon efficient FLIM instrumentation, the technique also becomes attractive to screen, with high temporal resolution, for fast changes in Förster Resonance Energy Transfer (FRET), such as those occurring upon activation of cell signaling.

We studied the effects of siRNA-mediated individual knockdown of an extensive set of 22 different phosphodiesterases (PDEs) on baseline levels and agonist-induced changes of the second messenger cAMP. Using HeLa cells stably expressing our FRET-FLIM sensor we imaged many hundreds of cells at 5 second intervals for each condition. Following segmentation of cells by the deep-learning implementation Cellpose, FLIM time traces were calculated and fitted for dynamic analysis with custom-made Python scripts. Taking advantage of the quantitative FLIM data, we found very limited effects of PDE knockdown on baseline and agonist-induced peak levels of cAMP. However, cAMP breakdown in the decay phase was significantly slower when PDE3A and, to a lesser amount, PDE10A were knocked down, identifying these isoforms as dominant in HeLa cells.

In conclusion, we present a robust platform that combines photon-efficient FLIM instrumentation with systematic gene knockdown and an automated open-source analysis pipeline. Our quantitative platform provides detailed kinetic analysis of cellular signals in individual cells with unprecedented throughput.

## Introduction

The genetic screens that have elucidated the roles of so many genes over the last decennia^1–3^ have, until recently, mostly relied on static end-point readouts such as cell viability or colony formation. However, in screens for genes involved in cell signaling and transient metabolic processes, static screening does not suffice, because the impact of such cell signals not only depends on the magnitude of the response, but also on its progression over time. Parameters like signal duration, inactivation, plateau phase and oscillatory behavior are essential for a complete understanding of signaling pathways. This necessitates genetic screens designed to capture the dynamics of signaling events, in short called “dynamic screens”. Thus, in recent years much effort has been invested into the development and improvement of methodology for dynamic screening^3,4^. Advanced live cell light microscopy is pivotal in these efforts, as it provides a large toolkit to address the dynamics of cellular processing in real time. These tools include time-lapse readout of single-cell morphological phenotypes by automated image analysis, and a variety of (fluorescent) indicators, both chemical dyes and genetically encoded indicators such as reporter constructs and FRET sensors. Combined with systematic genetic perturbations, imaging-based dynamic screening presents the state of the art in quantitative cell biology at the single cell level.

Our lab has focused on FRET because it can be used to study almost every aspect of cell signaling. FRET is a powerful, time-proven technique to study dynamic protein-protein interactions and also a great readout for biosensors, which can be designed to study various steps of signal transduction cascades. Consequently, biosensors have been widely adopted by the scientific community since they first became available in the early 2000s. For dynamic purposes, FRET is commonly detected either by ratiometry, in which the ratio of intensities of the FRET donor and acceptor are used to follow interactions, or by recording the donor fluorescence lifetime, i.e., the average time the donor stays in the excited state before returning to the ground state. While it is not easy to quantify FRET by ratiometry very precisely^5^, FLIM is a robust and inherently quantitative method for FRET detection which requires no additional calibrations or correction parameters^6^: interaction between donor and acceptor shortens the excited-state lifetime and is linearly related to FRET efficiency.

Thus, FLIM is ideally suited to quantitatively study baseline and stimulated FRET values in individual cells and among different cell populations, yielding data that are directly comparable between different laboratories around the world. The flip side of the coin is that dedicated and highly complex hardware is needed to read out the sub-nanosecond differences in donor lifetime that are associated with FRET. Moreover, the first implementations of FLIM equipment were custom-made, photon-inefficient and too slow for fast dynamic events. Therefore, until recently FLIM recordings were the exclusive domain of a few expert laboratories. In particular, the low throughput of FLIM rendered it ineffective for dynamic FLIM screens. However, over the last decade researchers have collaborated with leading microscopy manufacturers to come up with commercial implementations that circumvent some of these issues. First, convenient turn-key instruments were developed that opened up FLIM to non-expert users. Second, several groups, including our own, have collaborated with industry to devise and implement much more photon efficient and faster instrumentation, both for confocal^7^ and wide field microscopy^8^. These improvements enable following large numbers of cells in real time, with high data content and minimal photodamage, and make FLIM a very attractive choice for FRET-based signaling studies. Moreover, these instruments should now enable using FLIM in dynamic screening applications.

In this feasibility study, we developed a dynamic imaging-based screen and automated analysis for monitoring the activity of proteins involved in cellular signal transduction. We focused on the well-characterized second messenger cyclic Adenosine Mono Phosphate (cAMP), an ubiquitous messenger that controls many different cellular processes, including cell differentiation, gene transcription, secretion and ion channel activity^9^. cAMP is formed upon activation of G protein-coupled receptors (GPCRs) that signal via Gα_s_ subunit. Gα_s_ triggers members of the Adenylate Cyclase (AC) protein family, a group of isozymes encoded by 10 different genes in mammals, which rapidly produce cAMP from cytosolic ATP^10^. cAMP then relays the signal to a set of effector proteins, which include Protein Kinase A (PKA), a hetero-tetramer consisting of 2 cAMP-binding regulatory subunits and 2 catalytic subunits. Other effectors include nucleotide-gated ion channels, exchange factors such as Epacs and Popeye domain containing proteins^11^. In addition, recent studies have provided compelling evidence that the time course as well as precise subcellular localization of cAMP increases plays a pivotal role in determining the outcome of the signaling cascade^12–14^. For example, PKA members are often anchored to specific target sites by a family of A-Kinase Anchoring Proteins (AKAPs), further fine-tuning the biological effects of cAMP. The extensive set of proteins involved in synthesis of and response to cAMP underscores the importance of this messenger.

The kinetic properties of signaling are determined by the balance of production and degradation of cAMP. Cellular breakdown of cAMP is catalyzed by a family of specialized enzymes, the phosphodiesterases (PDEs). Based on their sequence relatedness, kinetics, modes of regulation, and pharmacological properties, the PDEs can be divided into 11 families^15^. In mammals, 3 of the 11 PDE families selectively hydrolyze cAMP (PDEs 4, 7, and 8), 3 families are selective for cGMP (PDEs 5, 6, and 9), and 5 families hydrolyze both cyclic nucleotides with varying efficiency (PDEs 1, 2, 3, 10, and 11). Selectivity in this case is defined as high substrate preference at physiological concentrations. Genes for individual PDEs can have multiple promoters, and the transcripts are subject to alternative splicing, resulting in nearly a hundred different PDE messenger RNAs^16^. However, most cell types express only a subset of PDE family members (e.g. in HeLa cells: PDEs 1A, 2A, 3A, 3B, 4A, 4B, 4D, 5A, 6A, 6C, 6D, 7A, 7B, 8A, 8B, 10A are expressed)^17^. In addition, the activity of PDEs can be further regulated at the protein level, for example by other second messengers including cGMP, Ca^2+^ and other PDE isoforms generating crosstalk between second messenger systems^18^ and further increasing the complexity of cAMP signaling.

In this study, we systematically investigated the breakdown efficacy of 22 different PDEs in HeLa cells by siRNA-mediated knockdown. siRNA-mediated knockdown has been proved to be an effective strategy to diminish PDE activity, as was shown by Willoughby et al, who focused on the role of PDE4 in HEK293 cells^19^. We created cell lines stably expressing the Epac-based cAMP FRET-FLIM sensor, Epac-S^H189^, which was generated in our lab by a series of sequential refinements^20,21^. Epac-S^H189^ consists of most of the sequence of Epac-1, with mutations to render it catalytically dead as well as dislodge it from membranes by deletion of the DEP domain. This moiety further harbors a single point mutation, Q270E, which increases its affinity for cAMP. EPAC is flanked by FRET donor mTurquoise-2 and a tandem of dark Venus proteins as acceptor^20^ which specifically tailors EPAC-S^H189^ for FLIM analysis. Upon cAMP binding, the Epac moiety undergoes a conformational shift, which decreases FRET and thereby increases the donor lifetime. The high FRET span and photostability of this sensor made it ideal for rapid screening purposes when the photon budget is limited.

In HeLa cells grown on 96-well plates, a specific PDE was suppressed in each well with a set of 4 different siRNA oligonucleotides, administered 72 h prior to imaging. For completeness of the feasibility study, we included all PDE families, irrespective of their selectiveness for cAMP. We monitored the production and breakdown of cAMP using a Leica SP8 FALCON microscope^7^ for high-throughput and photon-efficient recording of donor fluorescence lifetimes. Cells were automatically segmented using an established deep-learning based segmentation protocol, Cellpose^22^, and the various kinetic properties of cAMP signals in the cell interior were extracted by custom-made Python analysis routines. The highly quantitative results of 6 independent screens identified two dominant PDE species in determining cAMP breakdown in HeLa cells.

## Materials & Methods

### Stable expression of Epac-S^H189^ biosensor

HeLa cervical cancer cells (ccl-2) were cultured in DMEM (Gibco, 31966-021) supplemented with 10% FCS (Gibco, 10270). For creation of the stable cell line expressing the Epac-S^H189^ biosensor^20^ transfection of HeLa cells was performed with the *Tol2* transposon system^23^. For transfection two plasmids are used: a cDNA with the transposase sequence and another cDNA with the following elements: *Tol2*, promoter, the puromycin resistance gene, gene encoding for Epac-S^H189^ and a second *Tol2* sequence.

HeLa cells were seeded on 6-well plates at approximately 10% density and transfected the next day. 1 μg of both plasmids was mixed with 6 μl FuGENE reagent (Promega E269A) in 200 μl serum free DMEM and incubated for 30 minutes before adding the transfection mix to the cells. The cells were further incubated for 48 h and subsequently subjected to puromycin selection (1 μg/ml, SIGMA P8833). After 4 days cells were sorted on a fluorescence-activated cell sorter (FACS) based on mTurq2 fluorescence intensity.

### Generating PDE knockdown cells

Individual PDE gene knockdown was achieved by transfection with a pool of four exogenous short RNA duplexes (Table S1) with Lipofectamine® RNAiMAX cationic lipid formulation (ThermoScientific, 13778030). After incubation for at least 48 h, cells were imaged in fresh serum-free F12 culture medium (Gibco, 21041-025) on 96 well cell culture microplates (Greiner Bio-one, 655090).

### Stimuli used in the screen

Isoproterenol, Propranolol, Forskolin, IBMX (3-isobutyl-1-methylxanthine) and Cilostamide were purchased from Sigma-Aldrich.

### FRET detection for monitoring dynamic changes in cellular cAMP levels

To monitor the production and breakdown of cAMP in real time, the donor (mTurquoise2) fluorescence lifetime of the Epac-S^H189^ FRET biosensor was measured by FLIM. This FLIM sensor features a tandem dark (i.e., non-emitting) Venus acceptor which allows recording a large part of the donor emission spectrum while minimizing contamination of the signal with acceptor emission^20^.

FLIM experiments were carried out using a Leica TCS-SP8 FALCON confocal FLIM microscope^7^. The microscope was equipped with a humidified incubator with 5% CO_2_ at 37°C. Cells were excited with a pulsed diode laser (PicoQuant) at 440 nm, and photon arrival times were recorded with two HyD detectors, together covering the mTurquoise emission spectrum, adjusted to count photons at approximately equal rates (460-510 nm and 515-600 nm, respectively). In an additional channel, cell nuclei stained with 1 μM SiR-DNA (Spirochrome, SC007) for at least 45 minutes were imaged at 640 nm excitation, 650-725 nm emission, to support segmentation.

The recorded photon arrival time histograms showed multi-exponential decay, indicating the superposition of different FRET states. These FLIM data are well described by a double-exponential fit: a high FRET state with a lifetime of 0.6 ns, and a low FRET state with a calculated lifetime of 3.4 ns. For the lifetime analysis, the images were binned (2×2 pixels) and the pixel photon arrival times were fitted to a double-exponential reconvolution function with fixed lifetimes at 0.6 ns and 3.4 ns using Leica LAS X software. The resulting two images contained the amplitudes of these two components and were saved as TIF files, reducing the amount of raw data more than 40-fold.

For stimulation by photo-release of caged cAMP, cells were treated with DMNB-caged cAMP (4,5-Dimethoxy-2-Nitrobenzyl Adenosine 3’,5’-Cyclicmonophosphate, Molecular Probes, D1037) for at least 30 minutes prior to imaging at a final concentration of 1 mM. Uncaging was with a 200 ms UV pulse from the Leica EL6000 lamp (4 mW, approximately 400 mW/cm^2^ in our experimental setup with the 20x, 0.75 NA dry objective) using a DAPI_LP filter cube (380/40 nm, 405 nm dichroic) which was inserted in the confocal excitation path to enable UV illumination and confocal FLIM recording simultaneously.

### Kinetic analysis using Python

A graphical representation of the automated workflow and analysis pipeline is shown in Fig 2. All raw data is available on Zenodo^24^ and custom written software with the link to the corresponding Zenodo data repository can be found on our GitHub page^25^. These are the steps that are taken in the analysis, they can be repeated by running the software found online:

1. Cells are segmented using the deep learning algorithm Cellpose^22^. Due to minimal cell movement during acquisition, intensity data from all frames can be combined. The mean intensity is sent to Cellpose for deep-learning based cell segmentation.
2. Since the data contains the intensity of two fitted lifetime components, we calculate the weighted lifetime per frame using formula: 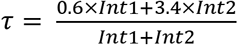, where τ is the average lifetime from the two fitted component amplitudes (*Int1* and *Int2*).
3. A lifetime trace is generated for each Region of Interest (ROI) in the labelmap, and fitted with the logistics function: 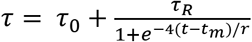, where *τ_0_* is the value of the baseline lifetime, τ_R_ is the range of lifetimes, *t_m_* is the time at the midpoint and *r* is the breakdown time. *r* represents the time required to break down the entire range if the breakdown had been constant at the value of the midpoint.
4. Goodness of fit was assessed by calculating the root-mean-square deviation (RMSD) and the corresponding mean absolute percentage error (MAPE). Only traces with MAPE below 1% were included in experiment evaluations and statistical analysis.
5. In the final step of the analysis is the generation of the figures using individual Jupyter notebooks that can be found in the GitHub repository^25^.

### Comparison of Cell segmentation

We initially used a conventional image analysis approach for cell segmentation by implementing Voronoi segmentation in an ImageJ macro (Github^24^), using the thresholded signal of nuclei stained with SiR-DNA as seeds. Each ROI created in this way was refined by restricting the ROIs with respect to size and intensity. Note that nuclear labeling proved not to be necessary for Cellpose segmentation. In comparing two independent segmentation runs, one with and the other without inclusion of the SiR-DNA channel, Cellpose yielded comparable high-precision labelmaps for all imaged FOVs. The results of Voronoi segmentation and Cellpose were in very good agreement, with the former providing significantly faster segmentation, at the expense of slightly reduced reliability and dependance on nuclear seeds. Since analysis time was not restrictive, we include segmentation with Cellpose in this study. For further details, see Fig. S1 and the text.

## Results

### Optimizing screening conditions and FLIM analysis

We first set out to determine optimal conditions to acquire time-lapse FLIM images using the Leica SP8 FALCON system. This system is designed for high-count rate Time-Correlated Single Photon Counting (TCSPC) and records mTurquoise2 lifetimes reliably at count rates in excess of 40 MHz per detector. We spread the mTurquoise2 emission over 2 HyD (hybrid detectors) by adjusting the spectrometer settings (see M&M), effectively doubling the maximum count rate. Global fitting indicated dominant lifetime components of 3.4 and 0.6 ns, indicating the superposition of two different FRET states. Saturation of the sensor with cAMP, as induced by the treatment of cells with the direct AC activator forskolin, changed the relative magnitude of the two populations but not their lifetimes. All time-lapse images were therefore fitted with a n-Exponential Reconvolution model using two fixed lifetime components of 3.4 and 0.6 ns, and the intensities resulting from these fits were exported as tiff files.

High signal to noise (S/N) ratio of lifetime measurements requires large numbers of photons to be collected per frame from each cell. However, possible photodamage, bleaching, and the necessary throughput set upper limits to the excitation power and acquisition time. To reliably resolve small differences in cAMP concentration, we aimed to achieve a lifetime repeatability (i.e., deviations of consecutive baseline readings in the integrated signal of each cell) of less than 50 ps RMS, even for dim cells. With the conditions detailed in M&M, actual observed RMS of ~25 ps, n=6500 cells, was achieved for most screens. As the lifetime span of the Epac-S^H189^ sensor ranges from ~2.0 ns in the resting state up to 3.3 ns when maximally saturated with cAMP, S/N ratio is thus better than 40:1. It can be seen in Fig. 1 that this was sufficient to clearly resolve cell-to-cell variability in response to addition of norepinephrine (NE), which activates β-adrenergic receptors in HeLa cells. This S/N ratio also suffices to resolve cell-to-cell variability in baseline lifetimes, and thus in resting cAMP concentrations. The lifetimes of FRET sensors at resting state appeared near-normally distributed (2.34 +/− 0.05 ns, mean +/− SD, n=154), Fig. 1E. Interestingly, a small percentage of cells with slightly increased cAMP levels were found (Fig. 1, see arrows). When imaged 2 days after culturing, these cells usually grouped together, suggesting clonal differences in baseline cAMP levels in WT HeLa cells. We also noticed that in most cells cAMP levels do not return to the initial resting values after transient stimulation with NE.

**Figure 1:**
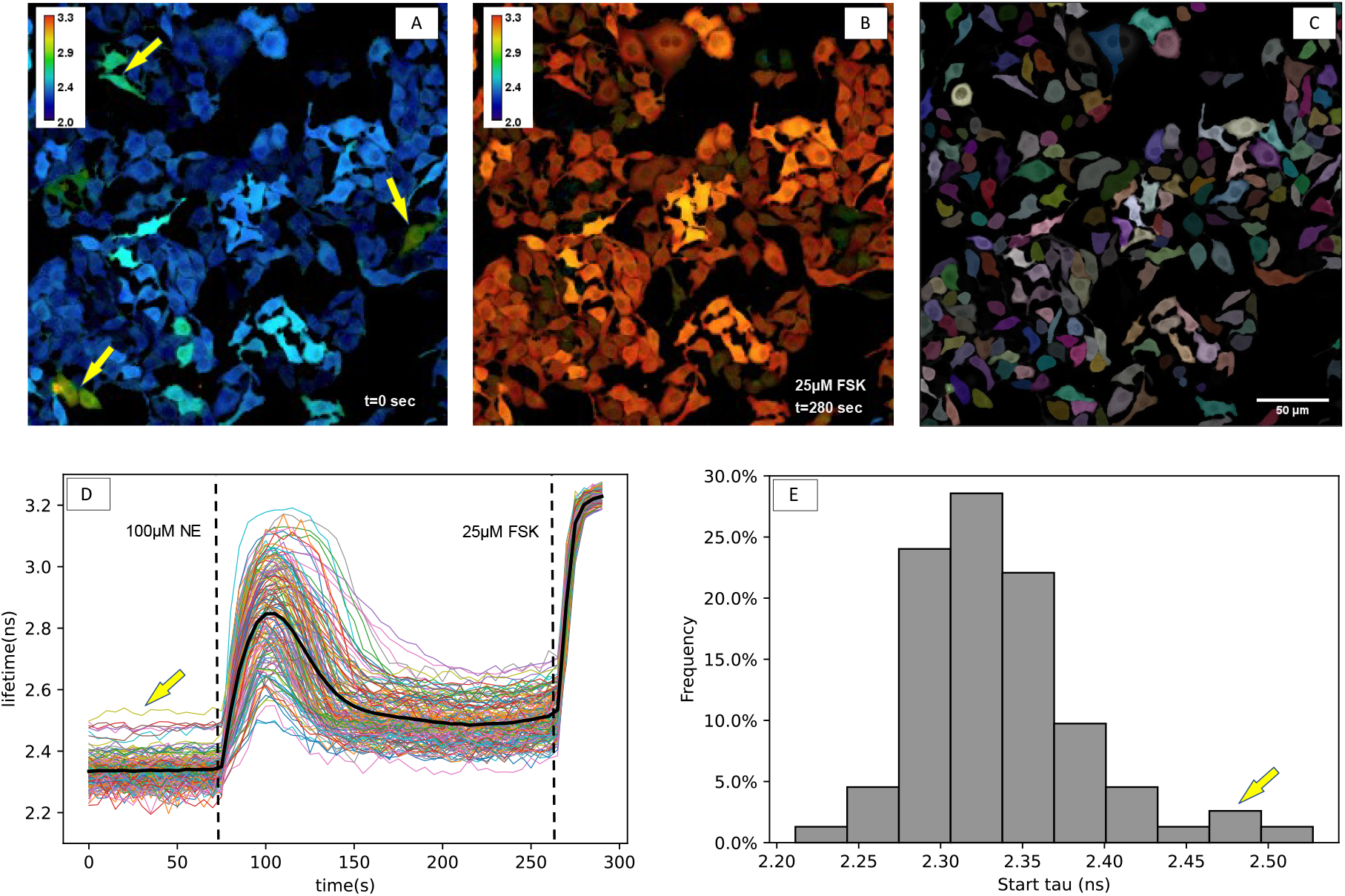
Detection of dynamic changes in cAMP levels in HeLa cells by FLIM. (A-C) Cells expressing the FRET-FLIM sensor Epac-S^H189^ are imaged at rest (A) and after stimulation with forskolin (FSK). (B) Calibration bar: lifetime in ns. Panel (C) shows the ROIs (color-coded) for each individual cell, as segmented using Cellpose, overlayed with fluorescence intensity. (D) Single cell FLIM time-lapse traces extracted from the same experiment show the transient response to stimulation with a 20-s pulse of 100 μM norepinephrine (NE). The bold black line represents the mean of all cells. NE was added at t=70 s and for calibration FSK was added at t=265 s. (E) Distribution of the baseline values (average of 20 samples for each cell). Yellow arrows in A, D and E indicate cells with higher baseline lifetimes.

**Figure 2:**
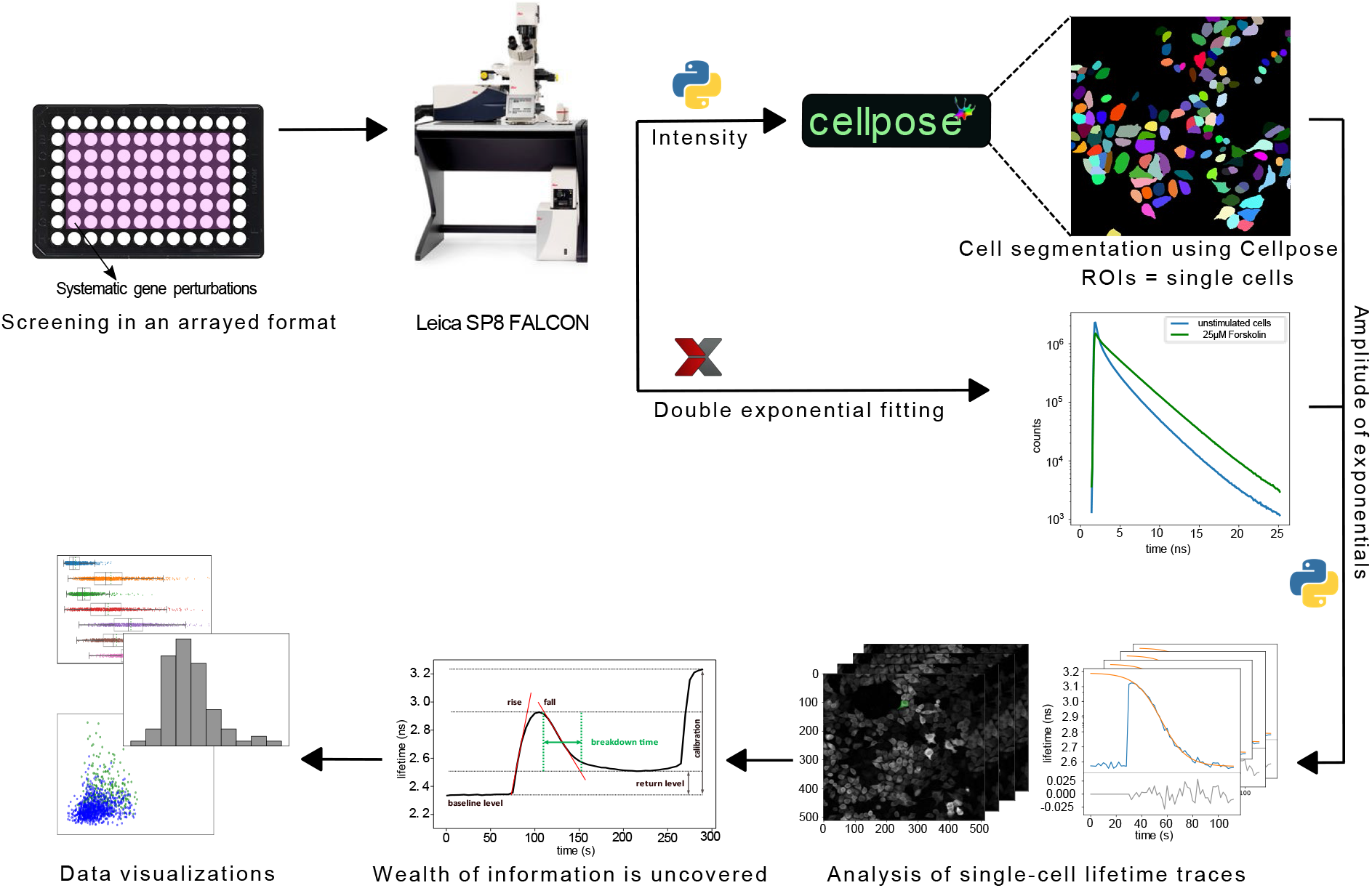
Schematic overview of the FLIM screen for dynamic changes in cAMP. HeLa cells expressing FRET-FLIM sensors grown in 96-well plates are treated with siRNA pools for 48 hours. Fluorescence was read out using an automated Leica SP8-FALCON FLIM microscope. The time-average of fluorescence intensity was used for segmentation using Cellpose, whereas the fluorescence lifetime data were fitted with a double-exponential decay using fixed fast and slow components of 0.6 ns and 3.4 ns, respectively. The magnitudes of those two components were exported to Python for further analysis. Based on the segmented ROIs, lifetime data were plotted for each individual cell, subjected to quality control, and agonist-induced changes were fitted with a suitable model. The fitting parameters are then summarized in descriptive plots.

The required throughput is determined by both the temporal resolution necessary to capture cAMP dynamics (time-lapse interval) and by the number of cells to be recorded from. The latter depends on several factors, including cell-to-cell variability due to stochastic differences inherent in signal transduction cascades and on incomplete penetration of the genetic perturbations carried out in the screen. Most siRNA mediated knockdown experiments display considerable variability in gene silencing resulting in incomplete or even no detectable knockdown in a percentage of cells^26^. Our pilot studies showed that recording from a few hundred cells in a single FOV captured most of the variation in each well. To minimize the risk that factors such as ongoing aging of the medium and increasing cell confluency might bias the results, we decided to run each entire screen, i.e. 22 knockdown conditions + controls in duplicate (60 wells in total), within 6-8 hours. Under these conditions, we found near-identical lifetimes in the experiments recorded at the onset and at the end of the 6-hour long screen (Fig. S2). Together, these considerations led us to conduct the screens using a 20x dry objective, recording a single field of view with ~200-600 cells per well, and at 2 or 5 second time-lapse rate.

### Automatic extraction of kinetic parameters

To optimize automated image analysis on a cell-by-cell base, we started by comparing algorithms for reliable segmentation of individual cells. We initially adapted standard image analysis methods by generating a dedicated Image J macro tailored to our cells. In essence, cell nuclei were detected by *in vivo* staining with SiR-DNA, followed by Voronoi segmentation to determine cell boundaries, which was based on the time-averaged intensity of the time-lapse images. This macro^25^ yielded good results, i.e. a ~95% reliable segmentation of cells was achieved as judged by eye. However, while our experiments were in progress, a general algorithm for segmentation of cells based on deep-learning algorithms was reported^22^, the performance of which we tested against our own developments. In several independent experiments we found Cellpose^22^ to be superior in reliability compared to more conventional image segmentation algorithms, including our own developments (Fig. S1). It must be mentioned that Cellpose is unpractically slow for near-real time analysis, but as it delivered very good segmentation without needing nuclear staining, we adopted it for all off-line segmentation of data in this study (for details, see M&M).

For each individual cell (ROI), we extracted mean fluorescence intensity and donor lifetime (Fig. 1) values, along with data on ROI size and potential error conditions such as disturbances by dislodged cells and out-of-boundary conditions (detailed description in accompanying information on our Github page). These data also were used to calculate RMS noise values of intensity and lifetime signals. Moreover, after fitting the agonist induced responses of cells to a suitable model (Fig. 2 and M&M), dynamic parameters such as activation rates, peak values, decay properties and steady state value were extracted. Next, we tested the reproducibility of our results with different batches of cells on different days. Our analysis showed excellent consistency of S/N ratio and calibration value following treatment with 25 μM forskolin. Baseline values were slightly more variable (Table S2), most likely reflecting small batch-to-batch variations in basal cAMP levels. These observations stress the importance of carrying out signaling screens within a limited time span, i.e., preferably on a single day.

### Caged-cAMP assay shows importance of PDE3A in regulation of cAMP breakdown

We next set out to conduct a FLIM screen to investigate the roles of the roles of individual PDEs in breaking down cAMP. We initially studied the kinetics of cAMP changes in HeLa cells upon photorelease of caged cAMP. For that, HeLa cells stably expressing the Epac-S^H189^ were seeded in 96-well plates, and using pools of 4 specific siRNAs against each isoform, individual PDEs were knocked down in duplicate wells. Cells were loaded with DMNB-caged-cAMP at 1 μM final concentration for 30 min. Uncaging with a 200 ms UV pulse caused an immediate increase in intracellular cAMP levels, and thus in donor lifetime, which subsequently returned towards its baseline level (Fig. 3). Hundreds of cells within a single FOV were imaged every 2 s for at least 140 s (or longer, if slow recovery called for that) and acquired data was stored for analysis offline. Following segmentation, time-lapse FLIM traces for each ROI were individually fitted to a logistic (sigmoid) curve. For large numbers of cells, the data and fits were visually inspected to ensure proper fitting using a Python script (results_browser25). cAMP breakdown time was then calculated from the resulting fit parameters and plotted for each PDE knockdown condition (Fig. 4). From these data, it is apparent that knockdown of PDE3A markedly affects the breakdown time in these cells (85.5 ± 2.5 sec, versus 37.9 ± 0.5 sec in WT cells, (mean ± SEM; p<0.001, student t-test). Additionally, a smaller but significant effect of PDE10A knockdown on cAMP breakdown was seen (51.4 ± 0.8 sec, p<0.001).

**Figure 3:**
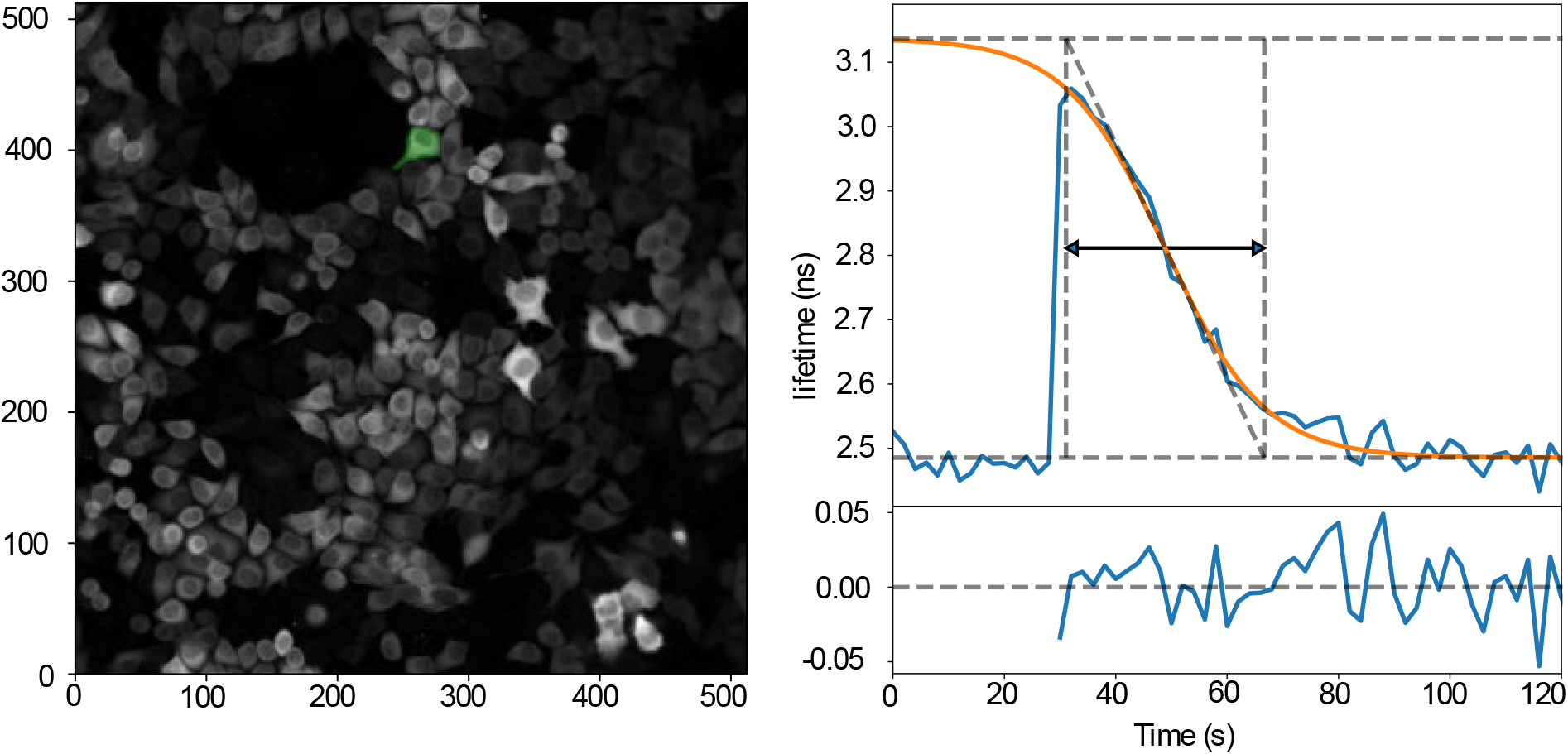
Changes in donor lifetime of the Epac-S^H189^ sensor upon uncaging of cAMP. The time trace (right) is from the green cell indicated in the left. Cells were imaged every 2 s and uncaging was at 25 s using a 200 ms flash of UV light. Note quick degradation of cAMP by PDEs back to baseline levels. Orange line shows the logistic function fitted to the data. Fit parameters are indicated by dashed lines: minimum and maximum lifetime (horizontal lines), maximum slope (diagonal line); vertical dashed lines indicate the intersection between maximum slope and min/max lifetime. The reported breakdown time (black arrow) is the time between the vertical two lines. The lower right panel shows the fit residuals.

**Figure 4:**
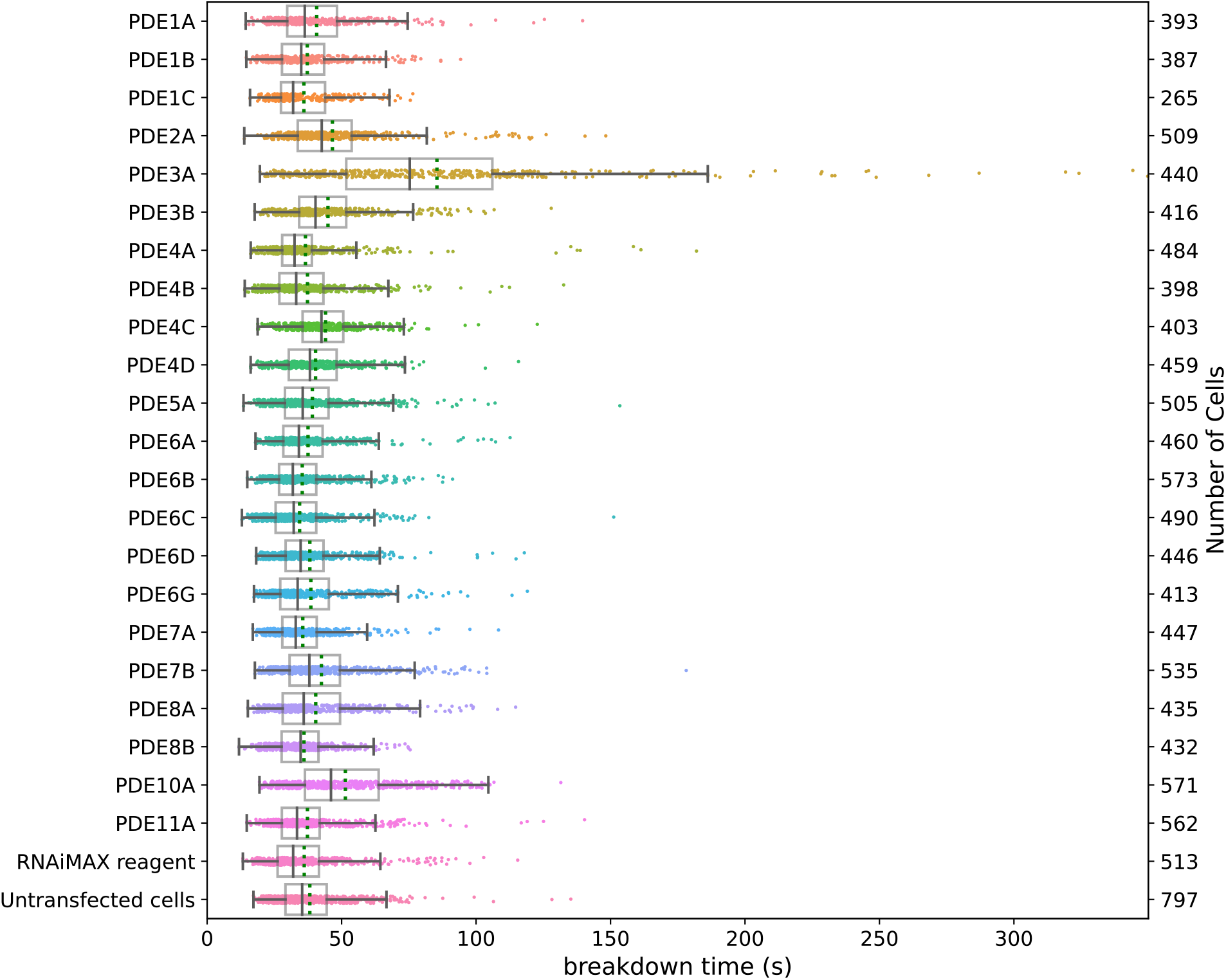
Decay time of donor lifetime signals following UV-uncaging of cAMP in cells treated with siRNAs for the indicated PDEs. Note significantly slower breakdown upon knockdown of PDE3A, and a smaller, but still significant contribution of PDE10A. Datapoints are fitted decay times of single cells. For each condition, the experiment was performed in duplicates, with cells grown, transfected, and assayed in two independent wells. Indicated are median value (vertical black line), mean value (green dotted line); boxes encompass middle 50% of values and whiskers represent 1.5 times the interquartile range.

From the data in Fig. 4, it is also apparent that the calculated decay times for individual cells show considerable variability. As the S/N ratio within a single cell is excellent (compare e.g. the variability of the initial 123 samples in the baseline in Fig. 4), the cell-to-cell variability in FLIM values likely has biological origin. Moreover, the extremely large span of the observations seen for knockdown of PDE3A and PDE10A suggest that lack of or incomplete PDE knockdown in individual cells is a further major determinant of variability in these wells. Furthermore, cell shape differences, e.g. in surface-to-volume ratios, are likely to affect cAMP clearance. This view is supported by the observation that very similar results were obtained when we repeated selected conditions, again in duplicate, a month later.

Next, we carried out a follow-up experiment to evaluate the effect of simultaneous knockdown of both PDE3A and PDE10A. Remarkably, knockdown of these two PDE genes in the same cell did not significantly slow down cAMP breakdown below the rate seen for PDE3A alone. This is perhaps unexpected because in PDE3A knockdown cells there is still a considerable rate of cAMP clearance. Therefore, next we assayed cAMP breakdown in cells pretreated with two well-characterized PDE inhibitors, the nonspecific PDE inhibitor IBMX^27^ (100 μM) and the PDE3 family specific inhibitor cilostamide^28^ (1 μM) administered either alone or together. Both inhibitors slowed down cAMP breakdown to rates slightly below that of combined PDE3A/PDE10A siRNA treatment, and combined they caused a further increase in cAMP clearance times (Fig. 5). It is also noteworthy that unlike PDE knockdown, inhibitor pretreatments selectively wiped out the population of cells with fastest breakdown times, consistent with the notion that high variability in breakdown speeds in the population of PDE3A and PDE10A knockdown cells reflects incomplete knockdown by siRNAs. Intriguingly, despite inhibition of all PDEs, cAMP still eventually is cleared in these HeLa cells. The mechanisms involved remain to be elucidated in further studies.

**Figure 5:**
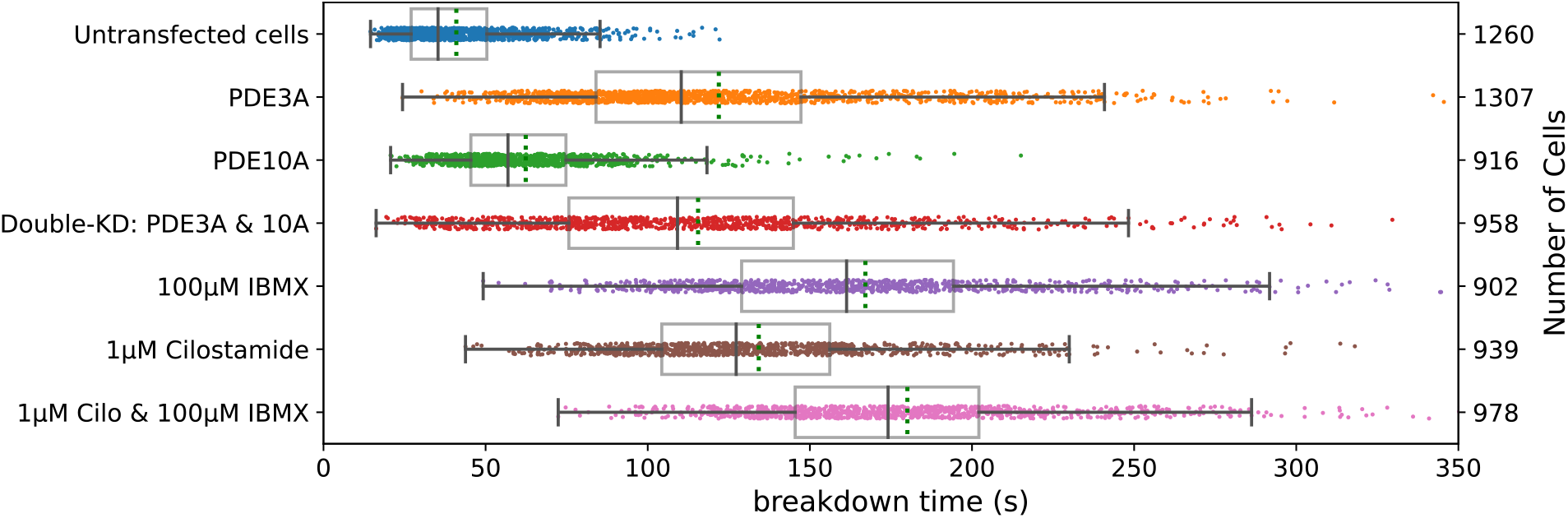
Breakdown of cellular cAMP after uncaging of DMNB-cAMP in HeLa cells. Datapoints represent the cAMP breakdown time values for all analyzed ROIs (individual cells) at a given condition. For each condition, the experiment was performed in duplicates, with cells grown, transfected, and assayed in two independent wells. Further details are as in Fig. 4.

While analyzing these data, we noted that baseline donor lifetimes in cells pretreated with DMNB-cAMP showed considerable biological variability, ranging between 2.4 ns and 2.7 ns (Fig. S3A). In contrast, untreated cells had average lifetimes of 2.28 ns at rest and showed considerably less variability (Fig S3B). The difference increased when cells were incubated with increasing concentrations of DMNB-cAMP, indicating some leakiness (spontaneous decomposition of the caging group in the cells) of this compound. In line with this, baseline donor lifetimes in PDE3A knockdown cells and in PDE3A / PDE10A knockdown cells were significantly elevated in DMNB-cAMP treated cells (Fig. S4A), but not in untreated controls. Together, these data indicate that PDE3A also has significant activity when cAMP levels are only slightly increased in these cells. We also noted that in the vast majority of stimulated cells, cAMP levels eventually returned to their pre-stimulation value (Fig. S4 C, D). A similar observation holds true for PDE knockdown cells.

We conclude that our screening platform is well suited to resolve even minor differences in kinetics of cAMP clearance kinetics, and that variability between experiments carried out several weeks apart is only minor. However, pretreatment with DMNB-cAMP appears to cause a significant disturbance of baseline cAMP levels, and this effect was amplified when PDE3A was knocked down. We therefore redesigned the experimental paradigm to circumvent the confounding effect of caged cAMP.

### Dominant role of PDE3A is confirmed using transient activation of GPCR signaling

Dynamic screens can also be carried out when AC is activated following stimulation of GPCRs with their cognate ligands. However, in such experiments it is much harder to dissect the contribution of PDEs in controlling the rate of cAMP clearance, because cAMP levels are also affected by the continued activity of proteins (GPCRs, G proteins and AC) upstream in the signaling cascade. Termination of Gα_s_ activity is believed to happen in seconds^29,30^ and AC activity is strictly dependent on GTP-loaded Gα_s_. However, receptor inactivation is much slower and, in most cases, not complete: a small proportion of receptors is thought to recycle to the plasma membrane where they can become reactivated by the agonist and continue to signal^31^.

Therefore, we adopted a protocol in which cells were stimulated with a receptor agonist, followed within 10-15 s by addition of excess of a potent competitive antagonist. We chose β-adrenergic receptors as they form a well-characterized G-protein coupled receptor system^32,33^, and are endogenously expressed in HeLa cells. We first stimulated HeLa cells with 40 nM isoproterenol which caused a rapid rise in cAMP levels and subsequently added propranolol at 60 nM concentration which caused a sharp decline following the stimulation. Finally, 25 μM forskolin was added to directly stimulate AC and obtain a maximal sensor response as a control. Fig. 6A shows a representative single-cell lifetime trace along with a fitted logistic curve capturing the decay kinetics.

**Figure 6:**
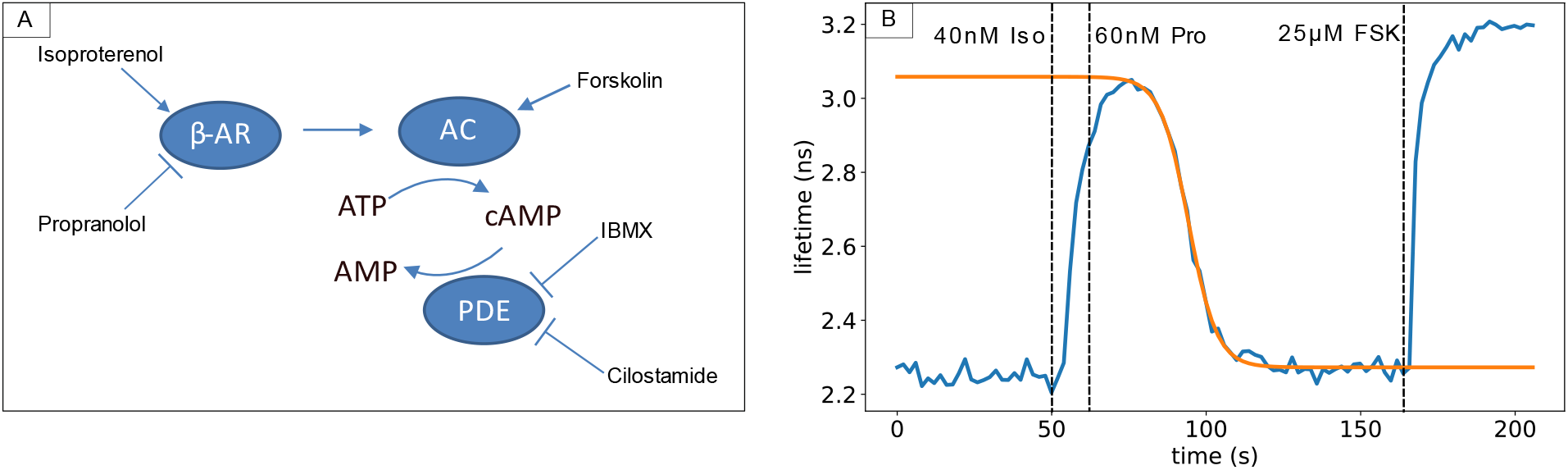
Assessing receptor mediated cAMP pathway. (A) Overview of the β-adrenergic receptor signaling pathway and agents used to affect cAMP production and breakdown. (B) Donor fluorescence lifetime changes of the Epac-S^H189^ sensor (blue line) in a single HeLa cell after stimulation of the β-adrenergic receptor with isoproterenol (40 nM) and subsequent blocking with the antagonist propranolol (60 nM). Forskolin (FSK) (25 μM) is added for calibration. Also shown is a logistic fit to estimate the cAMP breakdown time (orange).

Importantly, we noted that cAMP decay rates as determined following this experimental protocol in WT cells were approximately equal to those measured after photorelease of caged cAMP. This implicates that following addition of propranolol, all upstream steps in the signaling cascade became inactivated within seconds. It was also confirmed that when propranolol was added as first stimulus, no detectable response followed upon subsequent addition of isoproterenol. We therefore conclude that this experimental paradigm is well suited to study the role of PDEs in cAMP breakdown.

Fig. 7 shows effects of individual knockdown of the same set of 22 PDEs assayed according to this protocol. Again, we find that PDE3A is the most prominent enzyme controlling cAMP breakdown in HeLa cells, followed by PDE10A. The effects of knockdown of other PDEs were not significant. Furthermore, the double knockdown of PDE isoforms 3A and 10A together is also in good agreement with the data from the first screen after photorelease of caged cAMP. Remarkably, however, the effects of PDE inhibitors IBMX and cilostamide appeared more pronounced as compared to the first screen. Eventually, in all cases cAMP levels returned towards baseline levels, indicating the activity of additional cAMP clearance mechanisms in these cells.

**Figure 7:**
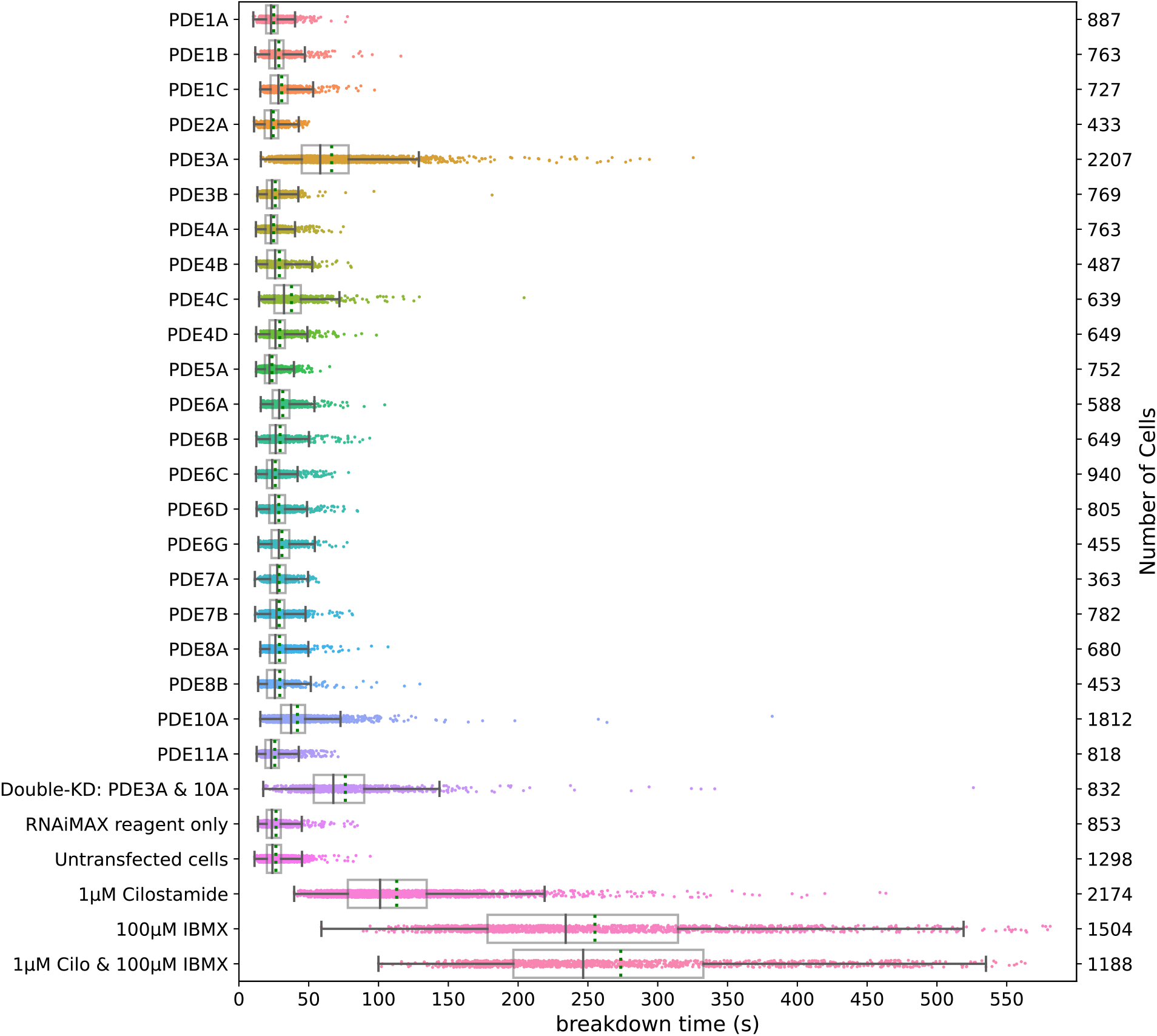
Breakdown of cAMP for different knockdowns of PDEs upon brief stimulation of the β-adrenergic pathway. For each condition, the experiment was performed in two independent wells. Further details are as in Fig. 4.

Finally, we tested for possible correlations between fluorescent properties of the cells, and outcome of the analysis. We found no evidence that cellular brightness affected donor baseline lifetimes for individual cells (Fig. S5). However, higher expression levels of the biosensor slightly prolonged cAMP breakdown, presumably due to the buffering effect of the Epac-sensor (Fig. S5). We conclude that variability in cellular cAMP breakdown speeds appears to dominated by true biological variability and that the agonist/antagonist stimulation paradigm is well suited to study the dynamics of cAMP turnover in genetic screens.

## Discussion

### Dynamic FLIM screening is finally within reach

The theoretical advantages of quantitative FLIM imaging for detection of FRET in screening applications have been long acknowledged: in principle, FLIM recording is insensitive to differences in expression levels of the sensor, sensor bleaching, excitation fluctuations and slight misfocusing. Moreover, because it is a single-channel technique, it also circumvents artifacts due to e.g chromatic aberration and sensitivity difference issues between channels which are often seen in ratio imaging. However, the practical implementation of FLIM in screenings has thus far been less than straightforward. This is because: first, until recently, available FLIM instrumentation has been either very slow or photon-hungry, and second, available FRET sensors most often are not optimized for FLIM read-out. In fact, FRET sensors have been almost exclusively optimized for ratio-imaging, and many sensors that offer a decent dynamic ratio change, show only a small lifetime change when assayed by FLIM^20,21^. Having spent significant efforts in improving both FLIM instrumentation and FRET-FLIM sensors, we here aimed to put these developments to the test and investigate the feasibility of a FLIM based dynamic screen. To this end we developed a screening platform and analysis pipeline to identify key PDE isoforms responsible for cAMP breakdown in HeLa cells.

We acquired data from HeLa cells expressing Epac-S^H189^ with a high-speed TCSPC instrument, the Leica SP8-FALCON and set up an analysis pipeline to extract dynamic data on cAMP levels in individual cells. Hundreds of cells per condition were segmented, and single-cell time-lapse traces with lifetime information were analyzed and fitted to a suitable mathematical model. We found that FLIM recording was stable and reproducible, with RMS noise levels in single-cell calculated lifetimes of about 25 ps, and that the majority of observed variability between cells and from day to day represents genuine biological variations. In addition, due to the excellent sensitivity of the instrument, no bleaching was detectable after collecting hundreds of frames from a FOV and we found no indications for phototoxic stress on the cells. Thus, FLIM recording has now become sufficiently fast and sensitive for routine data acquisition in single cell time-lapse screening experiments.

### Setting up the proof-of-principle screen

In order to characterize the roles of individual PDEs in HeLa cells, we employed two independent manners to induce transient increases in cytosolic cAMP levels. In initial screens, we uncaged DMNB-caged cAMP using a brief flash of UV light. The ensuing increase in sensor lifetime rapidly decayed towards baseline, and speed differences in cAMP clearance reflected the effect of knockdown of specific PDEs and PDE inhibitors. Among 22 individual PDEs included in our screen, we found PDE3A and PDE10A knockdowns to significantly slow down cAMP clearance. However, loading cells with DMNB-cAMP caused an increase in baseline levels, most likely due to some leakiness (spontaneous hydrolysis) of the compound, potentially biasing the results.

Therefore, we also tailored a protocol involving transient agonist-induced activation of GPCR signaling for screening purposes. GPCR signaling cascades encompass many different proteins at the cell membrane as well as in the cytosol, and cAMP metabolism following elevations due to receptor activation is highly complex. Upon stimulation with GPCR agonists, the time course of cAMP elevation is a mixture of AC activation and cAMP production on one hand, and cAMP clearance by PDEs on the other hand. Thus, the time course reflects direct aspects of Gα_s_ activity, but also indirect aspects such as GPCR inactivation (desensitization) and crosstalk with additional signaling cascades that may become activated downstream of these receptors. In addition, spatial aspects of cAMP signaling are critical for cellular function^34^ and they may affect the kinetics of lifetime changes in our screen. Lefkowitz et al. showed a putative switching mechanism at β2-adrenergic receptors, where protein kinase A (PKA) activated by the Gα_s_-pathway subsequently phosphorylates the receptor and causes enhanced Gα_i_ activation to limit further cAMP production, thereby forming a negative feedback loop^35^. In summary, it is not *a priori* clear that the outcome of a PDE-knockdown screen using GPCR activation should be one-to-one comparable to that of a cAMP uncaging screen. In an effort to focus on the role of PDEs in clearing cAMP from the cells, we adopted a protocol of transient GPCR activity using sequential agonist and antagonist stimulation. We stimulated β-adrenergic receptors, which are at the heart of a well-characterized GPCR system^32,33^, and abrogated signaling with the selective antagonist propranolol within 15 s. Interestingly, cAMP clearance rates following stimulation by this protocol appeared equally high as those following cAMP uncaging, indicating that Gα_s_ signaling and AC activity cease within seconds following receptor inactivation.

### Data fitting

In both designs of the screen, decay data were fitted with a sigmoid curve for pragmatic reasons. It may be argued that decay kinetics, certainly after cAMP uncaging, would approximate an exponential decay model if cAMP clearance is dominated by PDEs. However, different PDEs with potentially different affinities and expression levels would contribute to clearance and such a model would not accommodate a single value for enzymatic V_max_. More importantly, our data do not represent cAMP concentrations, but rather its binding to the Epac sensor, which itself is saturable. Finally, the observed cAMP breakdown curve is affected by the displacement kinetics of the agonist by propranolol. We therefore standardized fitting to a basic sigmoidal decay model. Fit residuals showed no systematic deviations, and Chi-squared values were around 1, indicating that the model validly describes the data.

### PDE3A and PDE10A are dominant in HeLa cells in both screening paradigms

In both screening methods, PDE3A and to a lesser extent PDE10A, were unequivocally identified out of 22 PDE isoforms as most important determinants of cAMP clearance. The relative importance of PDE3A has also been reported in several other cell types^36–38^, and thus this particular PDE isoform has become a potential drug target, most importantly in treatment of cardiovascular diseases and infertility^39–41^. Along with Cilostazol, an FDA-approved inhibitor for treatment of acute heart failure, alternative PDE3 inhibitors are currently being developed^42,43^. Also, several natural mechanisms by which PDE activity can be activated are currently under investigation for pharmacological manipulation^42^. Knocking down PDE10A alone also slowed down cAMP breakdown slightly but significantly. Nevertheless, rapid cAMP clearance was still observed both in cells treated with siRNA against PDE3A and PDE10A alone, or in combination. This indicates that other clearance pathways must be active. This view is supported by control experiments performed with the nonspecific PDE inhibitor IBMX^27^ and the PDE3-family specific inhibitor cilostamide^28^, which slowed down cAMP clearance to levels beyond those observed for the combined knockdown of PDE3A and PDE10A. It is intriguing that even in the presence of these two inhibitors together, cAMP levels still eventually decay towards baseline, suggesting the existence of additional, perhaps PDE-independent, clearance mechanisms.

### Limitations of the screen

Two important caveats must be made when interpreting these results. First, our sensor reads out cytosolic cAMP and we thus focused on the dynamics of cAMP generation and degradation throughout entire cells. Note that the readout of average cytosolic cAMP levels does not recapitulate the complexity of localized cAMP signaling, which over the last 15 years has been studied by several groups. For example, the Zaccolo lab has systematically addressed roles of specific PDE isoforms and protein phosphatases in creating and regulating local pools of cAMP^12,44,45^, unveiling e.g. the role of nanoscopic heterogeneity in cAMP signals in optimized cardiac contractility upon adrenergic activation^12^. In addition, a recent study by Lohse et al. demonstrates that under basal conditions, a large pool of cAMP in cells is bound, resulting in low free cAMP concentrations^46^. Nanometer-sized domains of even lower cAMP levels may be created and maintained by individual PDEs. GPCR stimuli may act to increase the concentration of cAMP so as to flood nanodomains and thereby trigger downstream effects. More generally, the compartmentalized signaling orchestrated by ACs and PDEs allows different GPCRs to generate unique spatially restricted cAMP pools that activate defined subsets of localized PKA, that in turn phosphorylate key targets in signal propagation, leading to specialized cellular responses. All of these aspects are not properly reflected in the results of the current study.

The second caveat is that the penetration of PDE knockdown by siRNAs is most likely far from optimal^26^. Although each gene was targeted by a pool of 4 different sequences, we have observed that control siRNA against lamin A provided marked depletion of this protein in up to 70% of cells, and even then, to varying degrees. Our screening data strongly suggest that PDE knockdown also has been variable, but this was not tested due to lack of availability of specific antibodies. In line with this interpretation, the effects seen with PDE inhibitors IBMX and cilostamide were more drawn out; for example, in Fig. 5 and 7 such pretreatment completely wiped out the fast-decaying population of cells. However, it is also conceivable that prolonged knockdown by siRNAs might have triggered compensatory mechanisms. For example, knockdown of PDE3A may cause other PDEs to partly take over its predominant role in cAMP breakdown. Such behavior would not be expected when cells are exposed to PDE inhibitors. Future studies using more state-of-the-art knockdown strategies such as the use of inducible degrons for depletion of PDE protein or CRISPR-Cas to completely knockout each PDE should address this point. The latter offers the added advantage that it is compatible with pooled microscopy screens, allowing for example addressing all PDEs in a single time-lapse experiment.

### Final conclusions

By combining high-end fluorescence lifetime imaging, a FLIM-optimized biosensor, deep-learning based cell segmentation and an automated analysis pipeline with systematic gene knockdown we achieved a robust screening platform for the systematic study of proteins affecting cellular signaling dynamics. Using open-source Python scripts and data structures, we illustrate the wealth of dynamic data delivered by quantitative time-lapse FLIM imaging. Whereas the multi-well format is particularly well suited for pharmacological characterization, including the analysis of effects of chemical libraries, the very quantitative nature of the obtained data should also be invaluable in extending such dynamic signaling screens to the pooled screening format. As such, we expect that this study will contribute to a substantial increase in throughput in signal transduction studies.

## Data availability

All data can be found on Zenodo repository: https://zenodo.org/record/4772516#.YKVO-6gzaUk Custom software can be found on GitHub: https://github.com/Jalink-lab/pde-screen-2021

## Author contributions

KJ, OK and SM conceived the study. OK and SM carried out microscopy screening experiments, and JK helped with cell culturing and establishing cell lines. RH and BB set up the cell segmentation and data analysis pipeline, and OK and KJ provided further additions to data analysis. The manuscript was written by KJ and OK, with input from all authors.

## Acknowledgements

We are grateful to the NKI Robotics and Screening Facility staff for help with siRNA plate preparation. We also acknowledge the input from members of the development team of Leica Microsystems, Mannheim. Dr Andre Nadler (Max Planck Institute of Molecular Cell Biology and Genetics, Dresden, Germany), prof. dr. Th. Gadella (University of Amsterdam, Amsterdam, The Netherlands) and prof. dr. H. Gerritsen (Utrecht University, Utrecht, The Netherlands) are acknowledged for advice and stimulating discussions.

