## Supplementary material for "Dynamic FRET-FLIM based screens of signal transduction pathways: a feasibility study": PDF of Supplemental Figs 1-4

***Supplementary Table S1. siRNA oligonucleotides used for individual PDE knockdown.*** *Gene knockdown was achieved by transfection with a pool of four exogenous short RNA oligonucleotides 72 h prior to imaging.*

| | | | | |
|---|---|---|---|---|
| PDE1A | GCACUAAGACGAUCAAAUA | UAAAUGGUCUUUCGAUGUA | CAAAUAGGUUUCAUCGAUU | GGAAGCAGUUUAUAUCGAU |
| PDE1B | GCAAGAAGAUGUGGAUUAA | GGAAGUACAAGAAUCCUUA | GUUCAGUGCUGGAGAAUCA | GAUGAGACACGGCAAAUCU |
| PDE1C | CCAAGGAGAUUGAAGAAUU | GAUCAUGCACUGAAAUUUA | CAUCAUCGCUGGACAAUGU | GAACUACUCACACGUUAUG |
| PDE5A | GAAGACAGCUCCAAUGACA | GAAAUCAGGUGCUGCUUGA | GAUGACAGCUUGUGAUCUU | GGAAACGGUGGGACAUUUA |
| PDE2A | GAACAUCCCUGACGCAUAU | CCAAUGAGAUGAUGAUGUA | GAGCUGAUCUACAAAGAAU | CAACAUCUUUGAUCAUUUC |
| PDE3A | GAAGAUAUCCCGGUGUUUA | CAAGGUAAAUGAUGAUGUU | UGACACAACUGCCAAACAA | GAGAUUGGAUAUAGGGAUA |
| PDE3B | GAAUACAACUUCCUUCUUC | GACAUCAAAUGCUGAAAUA | GAACAGCAAACAAAUAUUG | GGUGAUAAGUGGCUAACAG |
| PDE4A | UCACACACCUGUCAGAAAU | CCAAGCCGCUGGAGCUGUA | GGACAACUGCGACAUCUUC | GUAACAGCCUGAACAACUC |
| PDE4B | GAAUGUAGCUUGAGUAAAU | GAACUUGCUUUGAUGUAUA | GGAACCAGGUGUCUGAAUA | GAAAGAGACCUCCUAAAGA |
| PDE4C | AGAGACAGCUUUAGCCAAA | CCUCACAGCUAUCAUAUUC | CCAACCAGUUUCUGAUUAA | GGGACGGGCCUGACAGAUU |
| PDE4D | GAAAUCAAGUGUCAGAGUU | GAACUUGCCUUGAUGUACA | CCAAGGAACUAGAAGAUGU | GAACGUGGCAUGGAGAUAA |
| PDE6D | UGACGACGAUCUUCUUGUA | CGUCUUAACUGGGAACGUU | CAAACUAAAUUGGAUGAAC | AUCCCUAACUCCACAAAUA |
| PDE7A | GCUAGGAGAUGUACGUGUA | GGAUAGAGGUGAUUUAUGC | GUACUUCCAUUUAGAUAUG | GCGUGGAGCUAUUUCCUAU |
| PDE7B | GAAAUCAGUCCUCUUUGUA | GGCGAAAUCUUGUUUGAGA | CAACAGGCAGAAUGAAUUU | GCUGGGAGAUAUACGACUA |
| PDE6G | GAAAGGCGUUCAAGGGUUU | GAACAGACAUCACAGUCAU | CGACAUCCCUGGAAUGGAA | CGACAGACCAGGCAGUUCA |
| PDE8A | CAGCAUUUAACGUCAUAUG | UGCAGCAUUUAACGUCAUA | GUGCAGCAUUUAACGUCAU | CGUCAUAUGUGAUAAUGUG |
| PDE8B | GAGAAUAGCAGCAUAAUUG | GGUUAUAGAUGCAAUAUUG | GAAGUUCGCUCCCAGUUCA | GCGAUGACCACGUGAUUCA |
| PDE10A | GAACUAAACAGCUAUAUAG | GUAAUUGGUUUGAUGAUGA | GAACAAGGAGUUAUAUUCA | GCAGAGGCCUUGCCAAACA |
| PDE11A | GAAGAUUACUUGAUGCGGA | CCGACUGGCUAAUAAAUAA | UGAAGGAGCUCCAUUUACU | GGGAAGAGCUACACCAAAA |
| PDE6A | CAAGAGAGAUGAAGAGAUU | GAACAGGAGUGGACACAGU | CCAAUAACCUCUACCAGAU | GAGUCUGGAUGGAUGAUUA |
| PDE6B | GCACAGAAAUUUGCAAUGG | ACAAGGAGUUCUCUCGUUU | CAUUUGACAUCUACGAAUU | CCGGGUGGCUCAUCAAGAA |
| PDE6C | CAACUGACCUGGCUUUAUA | GAAAGGACCUGUAGACGAA | GGAUUUAUCUGUAACAUGA | CAGCAGAACUGUACGAAUU |

***Supplementary Table S2. Comparison of mean donor baseline lifetimes in unstimulated control wells.*** *Mean baseline values are calculated from all of segmented individual cells from the control wells of respective experimental day.*

| | mean baseline ± SD, ns | number of cells | max. mean baseline value |
|---|---|---|---|
| Exp. day 20190502 | 2.30 ns ± 0.08 | 305 | 2.55 |
| Exp. day 20191107 | 2.24 ns ± 0.04 | 1252 | 2.44 |
| Exp. day 20200107 | 2.32 ns ± 0.04 | 135 | 2.49 |
| Exp. day 20200206 | 2.29 ns ± 0.04 | 749 | 2.47 |

***Supplementary Figure S1. Comparison of cell segmentation routines: Cellpose versus custom Voronoi-based Fiji routine.*** *Images show segmentation outcome (labelmaps, in which each cell ROI corresponds to a color label) overlayed on the intensity image. The Voronoi-based segmentation generates cell labels separated by a 1-pixel border, which in general appear somewhat more 'blocky' and occasionally include the cytosol (edges) of neighboring cells. The script is designed to remove very dim and/or very small cells. Cellpose produces more smooth labels that better follow the outlines of the cells, but for dim cells often produces labels that are too small. In our analysis, these cells are removed in later analysis steps.*

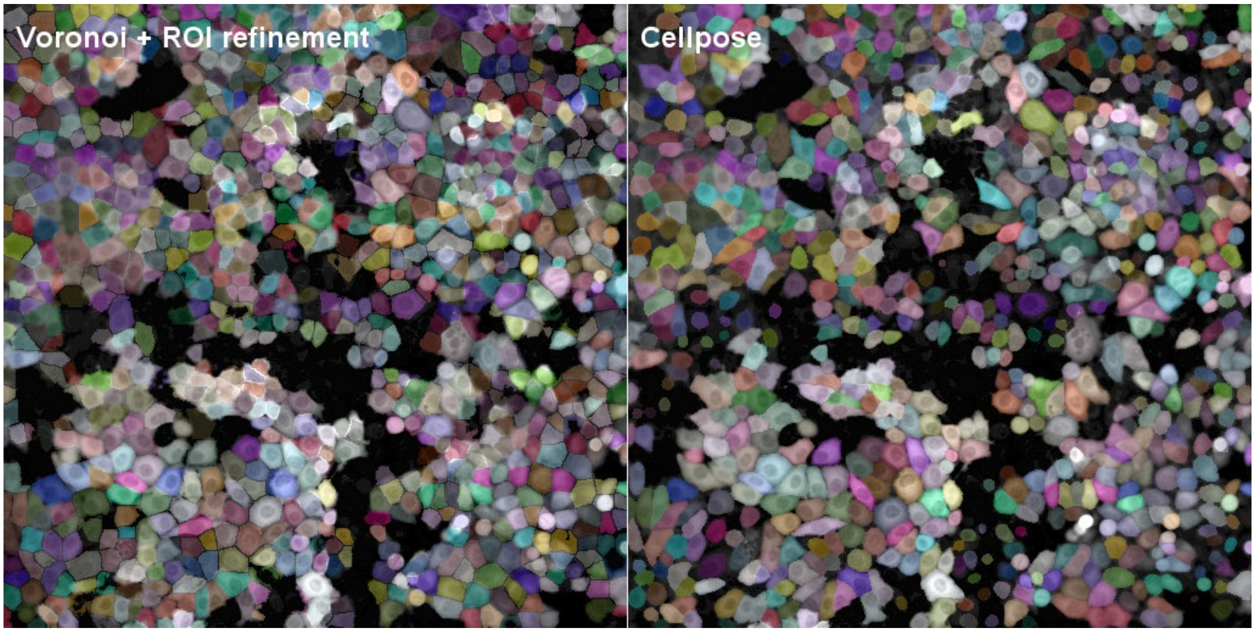

***Supplementary Figure S2: Baseline lifetimes, and thus cAMP levels, are stable for the duration of the experiment.*** *Distribution of baseline lifetimes of all cells in a well acquired at the onset of the screen (green, well B08), and a well acquired 6 hours later towards the end of the screen (blue, well G06). Well B08: 2.27 ns +/- 0.04 ns (mean +/- SD), max 2.44 ns, SEM = 0.00303, N = 210; well G06: 2.26 ns +/- 0.03 ns (mean +/- SD), max 2.39 ns, SEM = 0.00183, N=335).*

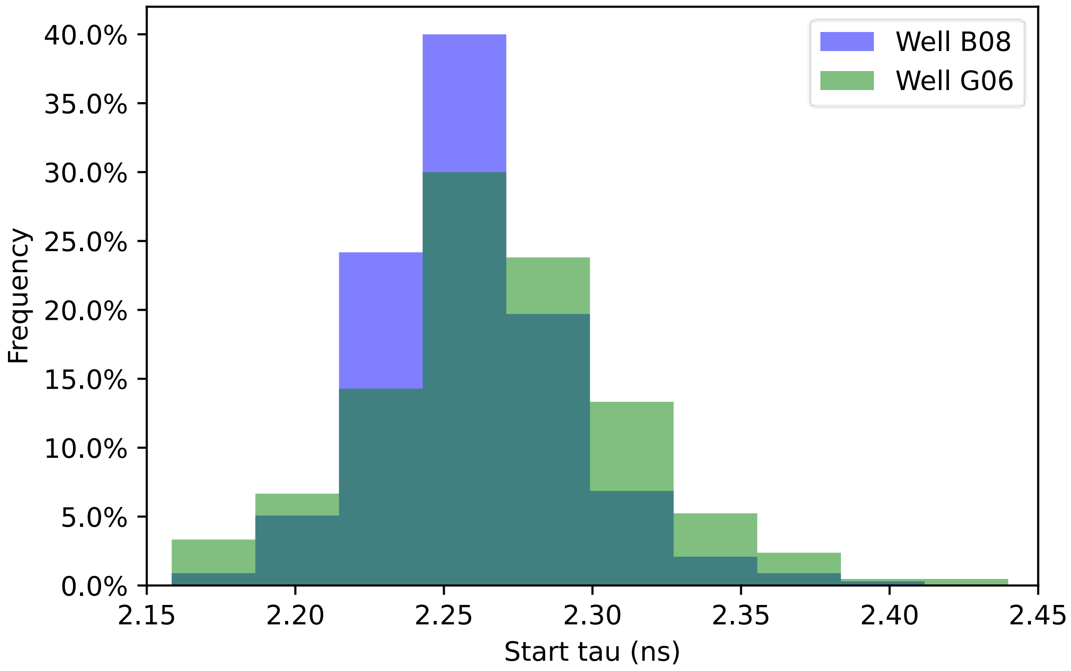

***Supplementary Figure S3: Analysis of base cAMP levels in both uncaging and receptor-activation screens.*** *(A) Lifetime traces of all cells in a non-transfected control well in a DMNB-cAMP uncaging experiment and (B) in an experiment with GPCR-stimulation. Each trace represents the intensity-weighted mean of all pixels in one ROI (segmented cell). A clear raise in pre-stimulation baseline levels is detectable in cells loaded with DMNB-cAMP. (C, D) Plotting 5 selected traces taken from panels A and B visualizes that upon breakdown, cAMP levels typically return to their pre-stimulation values in both types of assays. (E, F) Plots of pre-stimulation values (average of 12 baseline samples) versus post-stimulation return values (average of last 8 samples in the trace) for all cells plotted in A and B. Note the clear correlation between return levels and baseline levels.*

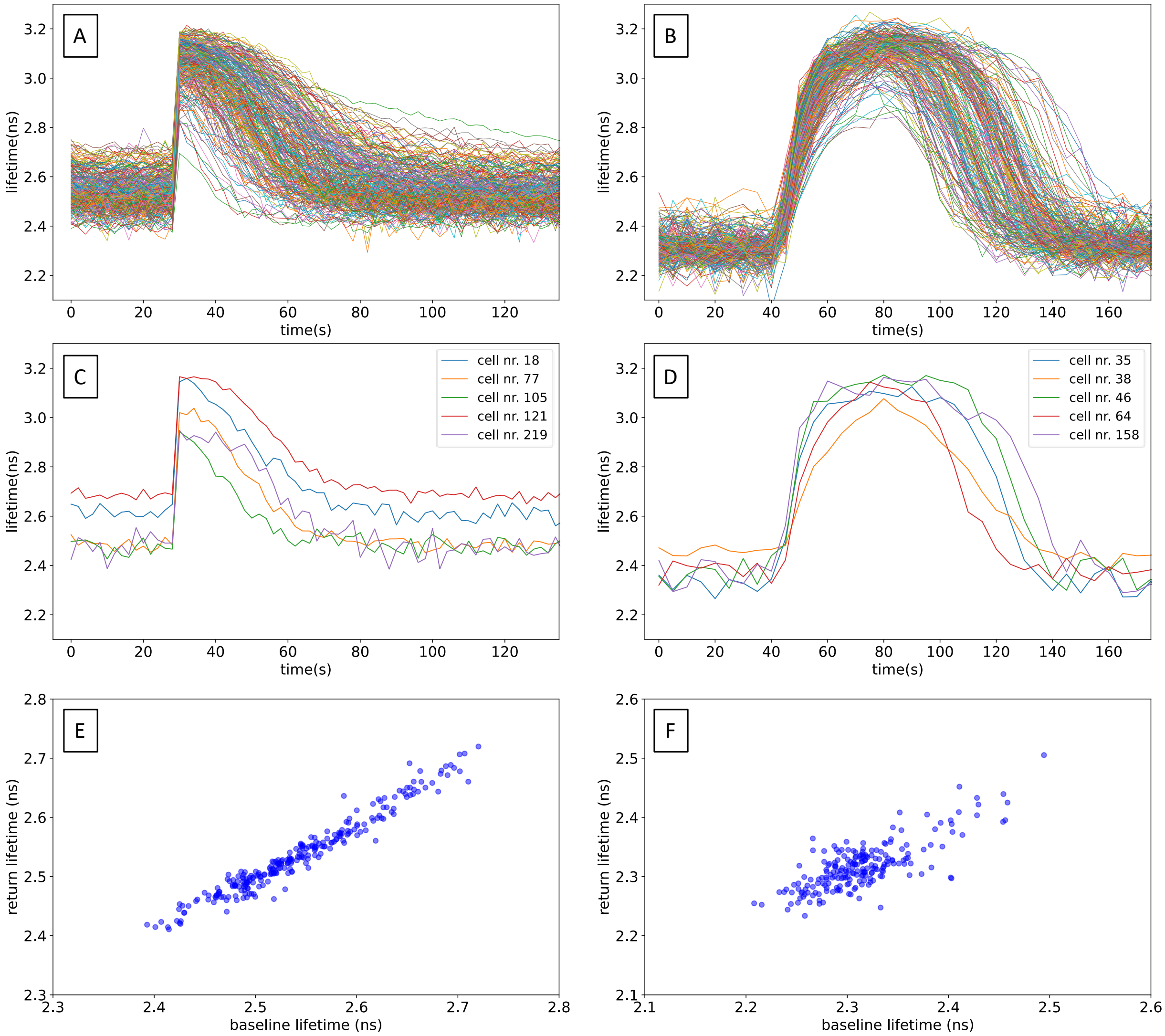

***Supplementary Figure S4: PDE3A and PDE10A knockdown slightly affects baseline cAMP levels in uncaging screens, but not in GPCR stimulation screens.*** *(A) Initial (baseline) donor lifetime values for all ROIs in the two screens with DMNB-cAMP uncaging and (B) in the two screens with GPCR stimulation (i.e., no loading with DMNB-cAMP). Note that PDE3A and PDE10A knockdown cells exhibit a slightly higher lifetime in cells pretreated with DMNB-cAMP, suggesting that those phosphodiesterases play a role in controlling baseline cAMP levels. In DMNB-cAMP loaded cells, treatment with PDE inhibitors also affects baseline lifetimes more prominently than in GPCR stimulation screens. (C, D) The difference between pre-stimulation lifetime values and the corresponding return lifetimes for all ROIs from two screens with DMNB-caged cAMP loading (C) and two screens without caged cAMP loading (D). Cells were treated with siRNAs for the indicated PDEs. Datapoints are fitted decay times of single cells. Each experiment was performed twice independently with duplicate wells for each condition (i.e., 4 wells per condition). Indicated are median value (vertical black line), mean value (green dotted line); boxes encompass middle 50% of values and whiskers represent 1.5 times the interquartile range.*

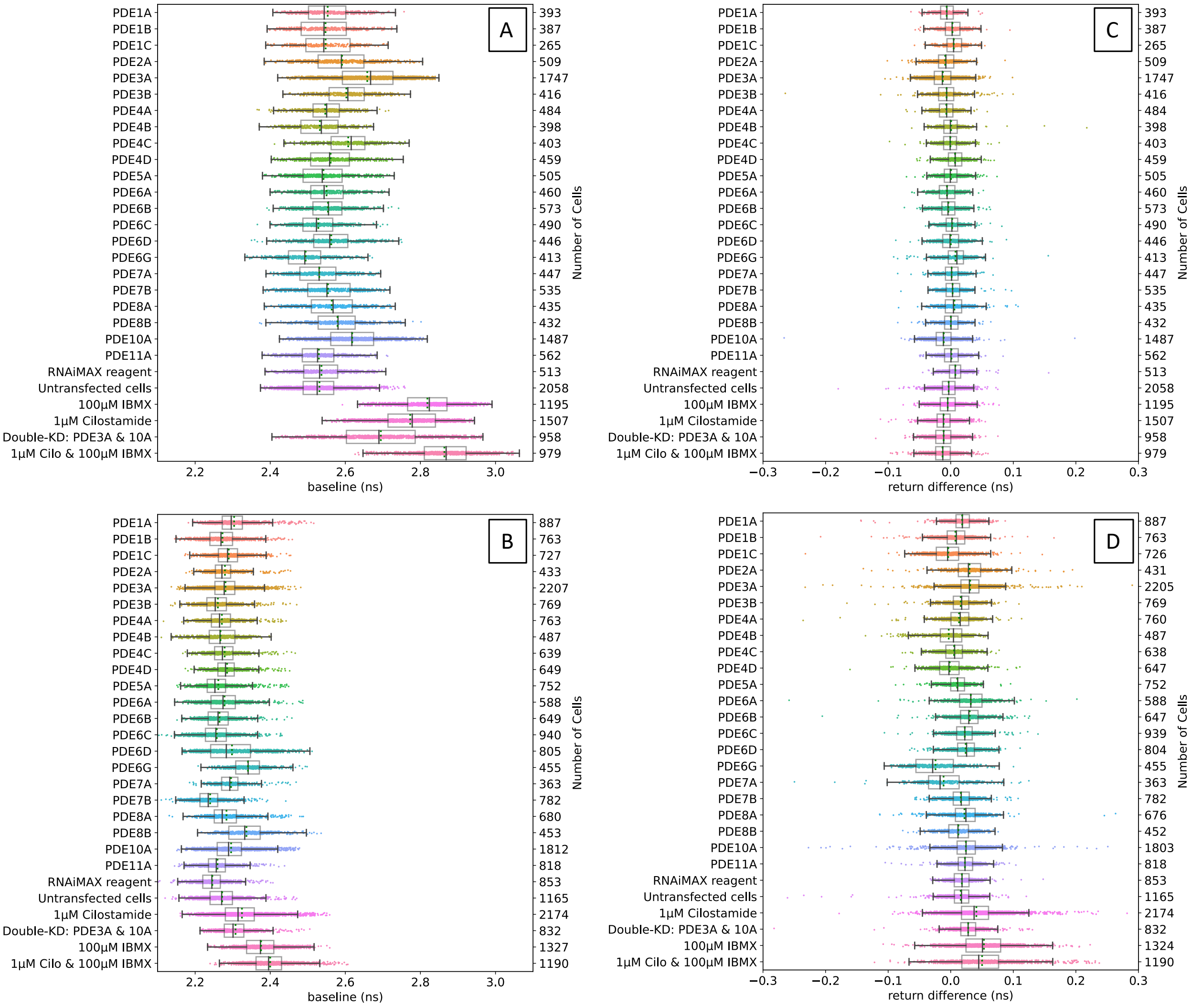

***Supplementary Figure S5: Relationship between biosensor expression levels, baseline levels and cAMP breakdown rates.*** *Initial (baseline) donor lifetime values are plotted against the corresponding cAMP breakdown times for cells expressing above-average (green dots) or below-average (blue dots) levels of bio-sensors. Data are pooled from the duplicate control wells (A, D), the PDE3A-knockdown wells (B, E) and from wells pretreated with 1 µM cilostamide (C, F) from one experiment using DMNB-cAMP uncaging (A, B, C) and one involving GPCR stimulation (D, E, F) carried out on two different days.*

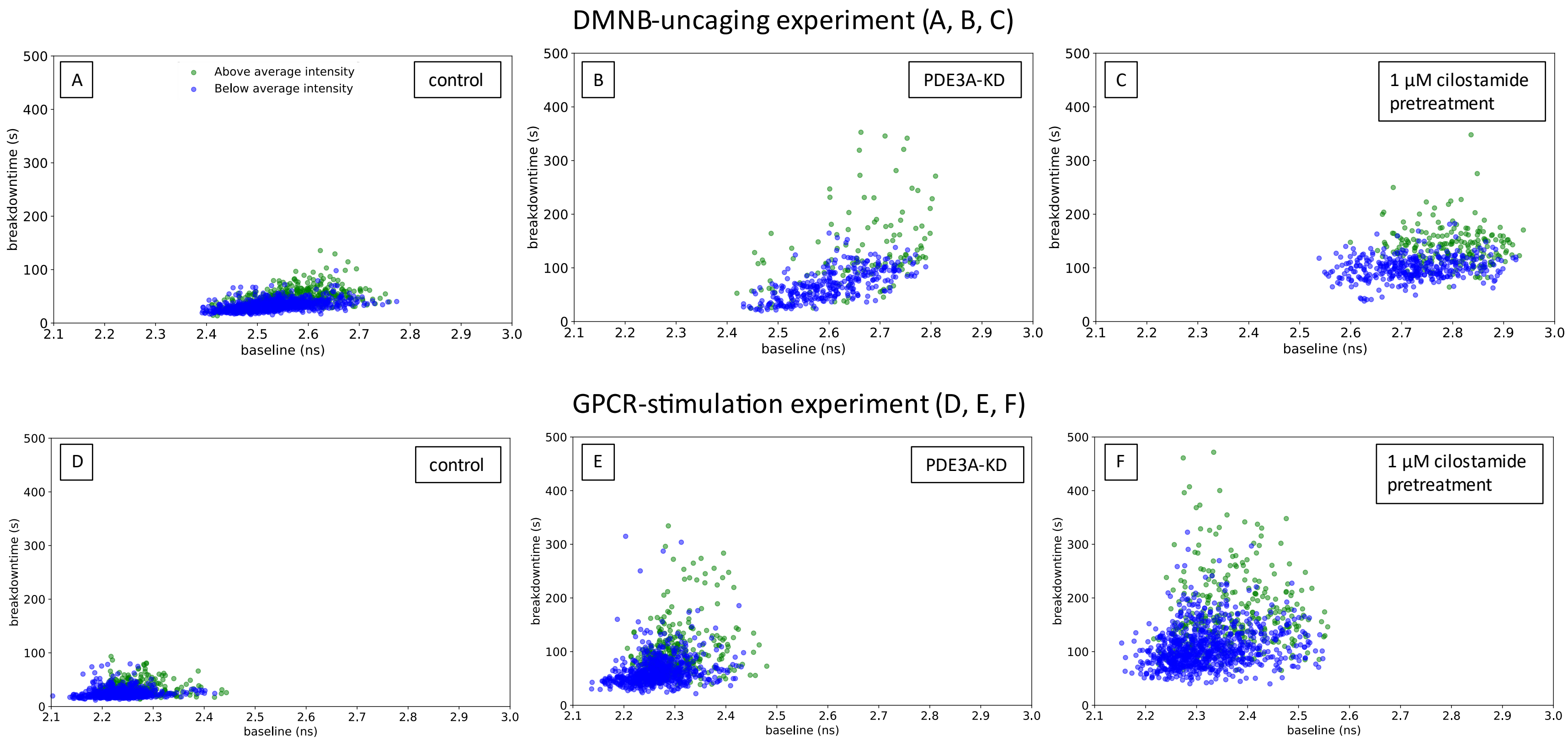
